# Evidence that the monoamine oxidase B (MAO-B) plays a central role in the inotropic dysfunction induced by genetic deletion of the Mas-related-G protein-coupled receptor D (MrgD) in mice

**DOI:** 10.1101/2024.03.27.586916

**Authors:** Lucas Rodrigues-Ribeiro, Julia Rezende-Ribeiro, Sérgio Scalzo, Maria Luiza Dias, Bruno de Lima Sanches, Marcos Eliezeck, Itamar Couto de Jesus, Joseph Albert Medeiros Evaristo, Kinulpe Honorato Sampaio, Diogo B. Peruchetti, Jader Santos Cruz, Fábio César Sousa Nogueira, Maria José Campagnole-Santos, Silvia Guatimosim, Robson Augusto Souza Santos, Thiago Verano-Braga

**Affiliations:** National Institute of Science and Technology in Nanobiopharmaceutics (INCT-Nanobiofar), Department of Physiology and Biophysics, Institute of Biological Sciences, Federal University of Minas Gerais, 31270901 Belo Horizonte-MG, Brazil; Department of Physiology and Biophysics, Institute of Biological Sciences, Federal University of Minas Gerais, 31270901 Belo Horizonte, Minas Gerais, Brazil; Proteomic Unit, Department of Biochemistry, Institute of Chemistry, Federal University of Rio de Janeiro, 21941909 Rio de Janeiro, RJ, Brazil; Laboratory of Proteomics/LADETEC, Institute of Chemistry, Federal University of Rio de Janeiro, 21941598 Rio de Janeiro-RJ, Brazil; Federal University of the Jequitinhonha and Mucuri Valleys (UFVJM), 39100000 Diamantina, Minas Gerais, Brazil; Department of Biochemistry and Immunology, Institute of Biological Sciences, Federal University of Minas Gerais, 31270901 Belo Horizonte-MG, Brazil; Department of Molecular Medicine, University of Southern Denmark, 5230 Odense M, Denmark

## Abstract

The renin-angiotensin system (RAS) plays a critical role in the regulation of the cardiovascular system. The Mas-related G protein receptor member D (MrgD) is the receptor of alamandine, and both are components of the RAS noncanonical arm. Alamandine/MrgD induces vasodilation, anti-inflammatory, anti-fibrotic and anti-oxidative effects. In contrast, *Mrgd* gene deletion leads to a remarkable dilated cardiomyopathy (DCM) in mice. Here, we aimed to investigate the molecular mechanisms of DCM triggered by the deletion of MrgD in the left ventricle and isolated ventricular cardiomyocytes from 8-12 weeks old mice using phosphoproteomics. Our findings revealed an increased oxidative stress not caused by angiotensin II/AT1 hyperactivation but instead due to the up-regulation of the monoamine oxidase B (MAO-B), leading to a higher catabolism of dopamine and epinephrine in the MrgD-KO cardiac tissues. The oxidative environment induced by MAO-B hyperactivation seems to be the cause of the observed alteration in ionic dynamics - altered Ca^2+^ transient and Na^+^/K^+^-ATPase activity - leading to altered resting membrane potential (RMP) and decreased contraction of MrgD-KO cardiomyocytes. In addition, cardiac Troponin-I phosphorylation, and Titin dephosphorylation seem to contribute to the contractile dysfunction observed in MrgD-KO. The treatment of cardiomyocytes from MrgD-KO mice with the MAO-B inhibitor Pargyline reverted the observed impaired contraction, corroborating the hypothesis that MAO-B hyperactivation is, at least partially, the cause of the failing heart observed in MrgD-KO mouse. The findings reported here provide important insights into the pathogenesis of heart failure and suggest a potential therapeutic target (MrgD activation) for managing failing hearts.

## INTRODUCTION

The renin-angiotensin system (RAS) plays a critical role in the regulation of the cardiovascular system, where it modulates the cardiac output, vascular resistance, blood volume, and blood pressure. RAS is composed of two functional axes acting most of the time in a counterregulatory fashion. The canonical axis - named in this way as it was identified first in the last century ^1^ - is composed mainly by angiotensin (ang) II and the receptor AT1. The non-canonical axis - also known as protective axis - is made of mainly by ang-(1-7), alamandine and their respective receptors MAS and MrgD. RAS impairment, tending for the hyperactivation of the ang II / AT1R axis, may precipitate inflammation, fibrosis, and hypertrophy in the heart while the opposite - ang-(1-7)/Mas or alamandine/MrgD activation - leads to cardioprotection ^2–4^.

Current data support a cardioprotective function for alamandine/MrgD axis. Alamandine and its receptor MrgD were first identified in 2013, in a publication that also reported that alamandine treatment reduced the cardiac fibrosis induced by isoproterenol in spontaneously hypertensive rats (SHR) ^5^. Later, another publication reported that the deletion of the *Mrgd* gene in C57BL6/J male mice (KO) precipitated a pronounced dilated cardiomyopathy (DCM), represented by contractile dysfunction, reduced ejection fraction, reduced fractional shortening, increased end-diastolic volume, increased end-systolic volume, and reduced cardiac output ^6^. However, the mechanisms by which *Mrgd* deletion leads to DCM are still elusive. Thus, we aimed to study the molecular mechanisms underlying the DCM induced by *Mrgd* gene deletion using proteomics, phosphoproteomics, and other analytical techniques. We observed a deleterious remodeling of the cardiac proteome and phosphoproteome in MrgD-KO male mice consistent with impaired redox system and excitation-contraction coupling mechanisms. Functional validation confirmed a significant oxidative stress in the Mrgd-KO cardiac tissue, increased MAO-B abundance and impaired contraction of MrgD-KO cardiomyocytes which was reverted by the MAO-B irreversible selective inhibitor Pargyline. In addition, we observed a differential phosphorylation in sarcomere proteins which correlate with impaired cardiac tension and the reported DCM phenotype of MrgD-KO mice ^6^. In summary, this work provides important cues of the molecular mechanisms underlying the association between the lack of alamandine/MrgD signaling and DCM.

## MATERIALS AND METHODS

### Materials

All reagents used in this study were acquired from the Sigma-Aldrich, except for the protease and phosphatase inhibitors (Roche), Collagenase type II (Worthington), and trypsin (Promega). Solutions were prepared using purified water from the Milli-Q system (Millipore). Low-binding microtubes (Eppendorf) were used for processing the samples.

### Animals

All animal care and experimental procedures were in accordance with the current edition of the NIH Guide for the Care and Use of Laboratory Animals. The experiments were approved by the local Animal Ethical Committee (CEUA-UFMG; protocol 5/2018). C57BL/6 (WT) and MrgD-KO (KO) male mice were provided by the BICBIO-2 animal facility from ICB-UFMG. The animals used in this study were 8-12 weeks old. For MrgD-KO, the following PCR primers were used for genotyping: MrgD-8, 5′-CATGAGATGCTCTATCCATTGGG-3′; reverse tetracycline transactivator primer (rtTA1), 5′-GGAGAAACAGTCAAAGTGCG-3′; and MrgD-1, 5′-CTGCTCATAGTCAACATTTCTGC-3′. All animals had free access to water and food. The temperature in the room was set at 23°C and a light-dark cycle of 12-12h was used.

### Isolation of left ventricle tissue

The left ventricle (LV) was obtained as published before ^7^. Briefly, the heart was dissected and immediately placed in ice cold PBS buffer solution pH = 7.4 containing (137 mM NaCl, 2.7 mM KCl, 8 mM Na_2_HPO_4_, 2 mM KH_2_PO_4_, protease and phosphatase inhibitors). Subsequently, the LV was rapidly dissected, snap-frozen in liquid nitrogen, and ground to powder using a mortar and pestle chilled with liquid nitrogen. After maceration, the samples were stored in ultra-freezer at −80°C for further use.

### Isolation of ventricular myocytes

The ventricular myocytes (cardiomyocytes; CM) were obtained as previously published ^8,9^. Briefly, the heart was rapidly removed and retro-perfused via the Langendorff method with modified Ca^2+^-free Tyrode solution (140 mM NaCl, 5 mM KCl, 5 mM HEPES, 1 mM MgCl_2_, 0.33 mM NaH_2_PO_4_, 10 mM glucose, and 100 U/mL insulin, at pH 7.4). Subsequently, the heart was perfused with Tyrode solution containing 50 μM CaCl_2_ and 1 mg/mL collagenase II for enzymatic digestion. Then, the tissue was mechanically dissociated and filtered to remove cellular debris. The extracellular Ca^2+^ ion concentration was gradually increased after three cycles of centrifugation and buffer exchange. Finally, the cells were centrifuged and kept in Tyrode solution. CM used for proteomic analysis were snap-frozen in liquid nitrogen and stored in ultra-freezer at −80°C. For patch clamp analysis, CM were used for experiments within 6 h after isolation.

### Western blotting

Proteins were separated by SDS-PAGE followed by western blotting. Primary antibodies and their sources are as follows: MAO-A (1:1500; Santa Cruz Biotechnology), ATP1A1 (1:100, Santa Cruz Biotechnology) and GAPDH (1:3000; Santa Cruz Biotechnology). Immunodetection was carried out using enhanced chemiluminescence detected with LAS 4000 equipment (GE HealthCare Life Science). Protein levels were expressed as a ratio of optical densities. GAPDH was used as a control for any variations in protein loading.

### Neurotransmitter measurements

We used a similar method as reported previously ^10^ for neurotransmitter measurements. Cardiac tissues were resuspended with a deproteinization solution made of 0.2 M perchloric acid and 3 mM cysteine. Subsequently, samples were sonicated in ice bath and centrifuged at 15000 x g for 15 min at 4°C. Supernatants were collected and analyzed using a Shimadzu HPLC system equipped with a fluorescence detector (HPLC-FD). A C18 chromatography column (250 x 4.60 mm, 5μ; Phenomenex) was used as stationary phase and a solution containing 12 mM CH_3_COONa, 0.26 mM EDTA.Na_2_ (pH = 3.5) was set as mobile phase. An isocratic gradient with 0.5 mL/min flow rate for 30 min was used. The following excitation (λ_ex_) and emission (λ_em_) wavelengths were used for the detection of monoamines: λ_ex_ = 279 nm and λ_em_ = 320 nm at high sensitivity and 4x gain. Protein amount from each sample was measured using a Nanodrop spectrophotometer (Thermo) to be used as a normalization factor. Standard solutions with concentration ranging from 0.006 to 1.536 ng/µL (0.006, 0.012, 0.024, 0.048, 0.096, 0.192, 0.384, 0.768 and 1.536 ng/μL) for Dopamine (DA), Epinephrine (E), Norepinephrine (NE) and Serotonin (5-HT) were prepared. The lower limit of quantification (LLQ) was determined considering the point where the peak was five times higher than the noise background (s/n = 5). The concentration of each neurotransmitter was calculated by plotting its measurements in a calibration curve followed by normalization based on the amount of the protein amount.

### Intracellular Ca^2+^ measurements

Intracellular calcium measurement was performed as previously described ^11^. Intracellular Ca^2+^ ([Ca^2+^]_i_) measurements were conducted using Fluo-4/AM (5 µM; Invitrogen)-loaded cells, incubated for 30 min. Following loading, the cells were washed with Tyrode solution to remove the dye excess. Subsequently, the cells were electrically stimulated at 1 Hz for 8 s to produce steady-state conditions, followed by a 6 s period of recording. All experiments were conducted at room temperature (23-25°C). Confocal line scan imaging was performed using a Zeiss LSM 880 confocal microscope equipped with an argon laser (488 nm) and a ×63 oil immersion objective located at the “Centro de Aquisição e Processamento de Imagens” (CAPI) at UFMG.

### Contractility acquisition

Following the isolation process, the myocytes were kept in Tyrode’s solution. Then, cells were subjected to electrical stimulation using platinum electrodes, with a pulse frequency of 1 Hz and a voltage of 30 V, each pulse lasting 5 ms. To capture cellular images (resolution of 640×480 pixels, featuring a pixel size of 0.25 μm/pixel and 8-bit depth), a high-speed digital CMOS camera (SILICON VIDEO 642 M, EPIX, Inc) was utilized, operating at a frame rate of 200 fps. The data from cellular contractility was obtained according to the protocol previously described ^12^. After basal state acquisition (Control), cells were treated for 10 min with Clorgyline (10^−5^ M) or Pargyline (10^−5^ M), a selective inhibitor of MAO-A or MAO-B, respectively, and contractility measurements were recorded again. Image processing and analysis were conducted using the CONTRACTIONWAVE software, employing the dense optical flow method ^13^.

### Patch clamp

Whole-cell recordings were obtained using an EPC-10 plus patch clamp amplifier (HEKA Electronics) at room temperature (23-25°C). To record action potential (AP), cardiomyocytes were maintained in external solution containing 140 mM NaCl, 5.4 mM KCl, 0.5 mM MgCl_2_.6H_2_O, 0.33 mM NaH_2_PO_4_, 11 mM glucose, 5 mM HEPES free acid and 1.8 mM CaCl_2_.2H_2_O (pH 7.4; adjusted with NaOH). Pipettes were filled with an internal solution containing 130 mM L-aspartic acid, 20 mM KCl, 5 mM NaCl, 2 mM MgCl_2_.6H_2_O, 10 mM HEPES free acid and 5 mM EDTA (pH 7.4). AP was evoked by applying a test pulse of 0.7 - 1 nA current for a duration of 3 - 5 ms at 1 Hz frequency.

### ATP measurement

The ATP content in LV or CM was measured using the ATP Assay Kit (Sigma-Aldrich; catalog number: MAK190) following the manufacturer’s instructions and using the Synergy HT spectrophotometer (Biotek). Protein quantification for data normalization was obtained using a nanodrop spectrophotometer.

### Sodium/potassium-transporting ATPase activity measurement

Na^+^/K^+^ ATPase activity was measured in LV and CM exactly as reported previously ^14^.

### ROS measurements

The dihydroethidium dye (DHE; Thermo; catalog number: D11347) was used to quantify the production of reactive oxygen species (ROS). Slides were equilibrated for 15 min with PBS at 37°C in a light-protected wet chamber. Transverse sections were incubated with DHE at a concentration of 10 μM for 30min at 37°C. The slides were washed three times with PBS and fixed with 4% paraformaldehyde (PFA) for 5 min and then sealed with mounting medium (glycerol 60%). Negative control sections received the same volume of PBS in the absence of DHE. Images were captured using the ApoTome.2 Microscope (Zeiss) using the laser at the 555 nm wavelength. Images were analyzed with ImageJ software (version 1.51j8). The integrative density of the fluorescence was measured and normalized by the selected area.

### Total collagen deposition analysis

Cardiac tissue was embedded in Optimal Cutting Temperature (OCT) (Tissue Tek) and cut into 8 μm thick using a cryostat (CM1850; Leica) with the temperature set as −20°C. cardiac tissue cross-sections were stained with the Picro-Sirius Red Stain Kit (Abcam; catalog number: ab245887) to stain total collagen deposition using the manufacturer’s suggested protocol. Images were captured using a camera connected to a light microscope with 40x magnification, and the Image ProPlus software (version 4.5.0.29) was used to quantify the total collagen deposition.

### Multiple reaction monitoring (MRM) for plasma RAS peptides quantification

RAS peptides were quantified in the plasma as described before ^15^. Briefly, venous blood was collected and inactivated using a cocktail containing protease inhibitors (Roche; catalog number: 04693124001). Blood samples were submitted to centrifugation and plasma collected. Plasma was cleaned up using a C18 solid-phase extraction (Sep-Pak, Waters) and reconstituted in 0.1% formic acid. Samples were analyzed by LC-MS/MS (Xevo TQ-S, Waters) using the multiple reaction monitoring (MRM) mode. For calibration curve, a stock solution containing the respective synthetic peptides (Bachem) was used. Data are presented as mean ± SEM and the parametric t-test was used for the statistical analyses.

### Sample preparation for proteomics and phosphoproteomics

Sample preparation was performed using the in-solution digestion protocol as previously reported ^7^. Approximately 10 mg of the cardiac tissue (grounded to powder) was resuspended in 100 μL of a lysis buffer containing 6 M urea, 2 M thiourea, 10 mM tris(2-carboxyethyl) phosphine hydrochloride (TCEP), 40 mM chloroacetamide, 50 mM triethylammonium bicarbonate (TEAB), and protease (Roche; catalog number: 04693124001) and phosphatase (Roche; catalog number: 04906845001) inhibitors. Samples were sonicated twice for 15 s under ice-cold conditions using a tip probe sonicator. Subsequently, they were incubated at 28°C in a ThermoMixer (Eppendorf) with agitation at 650 rpm for 2 h to allow reduction of disulfide bounds and alkylation of thiol groups by TCEP and chloroacetamide, respectively. Protein contents were quantified by the Qubit fluorometric method (Thermo). Solutions were diluted to a final concentration of 0.6 M urea, and 350 μg of proteins were digested using trypsin at a 1:50 ratio [enzyme (in μg) / sample protein amount (in μg) ratio] at 28°C for 16 h. Reaction was quenched by adding formic acid to a final concentration of 5% (v/v). Digestion efficiency was checked by mass spectrometry (Autoflex III MALDI-TOF; Bruker) at the CELAM, UFMG.

### Peptide labeling

Peptides were labeled according to the on-column dimethyl labeling protocol ^16^ with minor modifications. Briefly, digested peptides were reconstituted in 1 mL of 5% (v/v) formic acid (CH_2_O_2_). SepPak columns (Waters) were activated with 100% acetonitrile (ACN), conditioned with solution A (0.6% (v/v) acetic acid), loaded with the samples, washed with solution A, and then labeled with the respective labeling reagent for 30 min. For isotopic labeling, we used 50 mM sodium phosphate buffer (pH 7.5), 0.6 M sodium cyanoborohydride (NaBH_3_CN), and 4% (v/v) solution containing the respective formaldehyde: CH_2_O (light) or CD_2_O (medium). Thus, WT (*Mrgd* ^+/+^) and KO (*Mrgd* ^−/−^) samples were labeled using the light formaldehyde and medium formaldehyde, respectively. SepPak columns containing the labeled peptides were washed with solution A and then eluted with a concentration gradient (ACN 20%, 50%, 80%, and 100% (v/v) + 0.6% (v/v) acetic acid). The samples were dried down using a speedvac and reconstituted in 0.1% (v/v) TFA. The samples were quantified using a fluorescence-based method (Qubit; Thermo). The labeling efficiency was checked using a MALDI-TOF mass spectrometer (Autoflex III; Bruker) at the CELAM, UFMG. Subsequently, the light and medium-labeled samples were combined in a 1:1 ratio (KO vs. WT) and then checked again by MALDI-TOF MS. Finally, the samples were evaporated using a speedvac and stored at −20°C until further use.

### Phosphopeptide enrichment

Phosphopeptides were enriched using the TiO_2_ method ^17^ with minor modifications. Samples were resuspended in a loading solution containing 80% (v/v) ACN, 5% (v/v) TFA, and 1 M glycolic acid. Samples were then incubated for 30 min with titanium dioxide (TiO_2_) beads at a ratio of 6:1 [sample (μg) / TiO_2_ (μg)] under vigorous agitation. The loading solution containing phosphopeptides not bound to the resin in the previous step was transferred to a second tube and incubated again with TiO_2_ beads at a ratio of 3:1 [sample (μg) / TiO_2_ (μg)] for 30 min. After incubation, TiO_2_ beads were combined and washed consecutively with (i) 100 μL of the loading solution, (ii) 100 μL of 80% (v/v) ACN / 2% (v/v) TFA, and (iii) ACN 20% (v/v) / 0.1% (v/v) TFA. The tubes containing TiO_2_ beads were centrifuged, and the supernatant was collected. Phosphopeptides were eluted with a solution containing 1.5% (v/v) ammonia / 20% ACN. Eluted peptides were dried down in a speedvac and stored in −20°C until further use.

### Sample pre-fractionation

Samples containing the total proteome were subjected to offline reverse-phase pre-fractionation at high pH using C18 cartridges (SepPak; Waters). SepPak columns were activated with 100% ACN and equilibrated with 20 mM ammonium formate (NH_2_HCO_2_) at pH 9.5. The samples were reconstituted in NH_2_HCO_2_ 20 mM (sample pH was adjusted to 9.5) and loaded onto the SepPak column and washed with NH_2_HCO_2_ 20 mM. The samples were eluted into 18 fractions and combined in a concatenated manner into 9 final fractions [expressed as % (v/v) of ACN]: Fraction (F) 1 = 7.5% + 30%; F2= 10% + 32.5%; F3= 12.5% + 35%; F4= 15% + 37.5%; F5= 17.5% + 40%; F6= 20% + 50%; F7= 22.5% + 60%; F8=25% + 70%; F9= 27.5% + 80%. Samples were evaporated using a speedvac and stored at −20°C until further use.

### LC-MS/MS analysis

The proteome and phosphoproteome samples were resuspended in 0.1% (v/v) formic acid (solvent A) and analyzed using an HPLC EASY-nLC II (Thermo) configured with a two-column system and coupled to a Q-exactive Plus mass spectrometer (Thermo). The pre-column was approximately 3 cm in length and with an internal diameter of 100 μm, packed with C18-AQ Reprosil-Pur resin with 5 μm particle size. The analytical column (20 cm x 75 μm) was packed with C18-AQ Reprosil-Pur resin with a 3 μm particle size. The chromatographic gradient was set as follows: (i) 5-45% solvent B (95% (v/v) ACN and 0.1% (v/v) formic acid) in 32 min; (ii) 45-95% solvent B in 8 min, at a flow rate of 300 nL/min. The mass spectrometer was operated in a positive polarity and DDA mode. Eluting peptides were analyzed in MS1 with a mass range of 350-1800 m/z. The ions were accumulated up to 3×10^6^ or for 100 ms, whichever occurred first (MS1 “AGC target settings”). The parent ions were resolved with 70,000 resolution at m/z 200. The 15 most intense ions (Top 15) were selected in the quadrupole with a 2 Th isolation window for higher energy collision dissociation (HCD) fragmentation with a normalized collision energy (NCE) of 30%. The injection time was set at 50 ms or 10^6^ ions accumulated, whichever occurred first (MS2 “AGC target settings”). The fragment ions were resolved with 17,500 resolution at m/z 200, and their precursor ions were included in the dynamic exclusion list for 45 s. Raw spectra (.raw) were visualized with Xcalibur v3.0 (Thermo).

### Database search

The raw spectra (.raw) were deconvoluted using the MaxQuant software (version 1.6.17.0). Spectra were searched against the FASTA file of *Mus musculus* downloaded in December 2020 (17,492 entries) from the Uniprot/Swissprot protein database. The search parameters included a tolerance of 20 ppm for the first search, and the main search with a tolerance of 4.5 ppm. Dimethyl labeling was configured by activating the modifications “Dimethyl-Lys0” and “Dimethyl-Nterm0” for light labeling, and “Dimethyl-Lys4” and “Dimethyl-Nterm4” for medium labeling. Trypsin was chosen as the enzyme allowing up to two missed cleavages. Carbamidomethyl (Cys) modification was selected as a fixed modification, and oxidation (Met) and acetylation (protein N-terminal) were set as variable modifications. In the phosphoproteome samples, the “phospho(STY)” modification was enabled as variable modification. The “match between runs” option was activated with a time range of 0.7 min and alignment within a range of 20 min. For peptide identification, a minimum of 1 unique peptide was used. A reverse decoy database was used to determine the false discovery rate (FDR) that was set to <1%.

### Statistical analysis

Bioinformatic analysis was performed with Perseus (version 1.6.13.0) ^18^ and DanteR ^19^ software. Protein abundances were transformed to log2 values, and the missing values were imputed by random draw from a normal distribution with a width of 0.3 and a downshift of 1.8 relative to the standard deviation of measured values. The log-transformed intensities were normalized by the column median. Phosphopeptide abundances were normalized with the correspondent protein ^20^ from the proteome dataset. The one-way ANOVA was used to calculate the *p*-values using “DanteR”. Up-regulated features were defined when *p*-value < 0.05 and log2-transformed fold-change (log-FC) > 0.26. Down-regulated features were defined when *p*-value < 0.05 and log-FC < −0.26. Principal component analysis (PCA) was created using the “tidyverse” and “factoextra” packages in R. Gene ontology and volcano plots were created using the “ggplot2” and “ggthemes” packages, also in R. Interaction network graphs for KEGG and REACTOME pathways were created using Cytoscape 3.8.2 with the ClueGO 2.5.7 application. Boxplot graphs were created using GraphPad Prism 8.

For all other methods used in this study the student’s t-test was used to determine significance values, except for the contractile experiments with MAOs inhibitors (6 experimental conditions) that the one-way ANOVA with Tukey’s post-test was used.

## RESULTS

### Proteome and phosphoproteome analysis of the cardiac tissue derived from MrgD-knockout mice

Aiming to have a better understanding of the molecular mechanisms underlying the dilated cardiomyopathy (DCM) precipitated by the deletion of the alamandine receptor (MrgD), we compared the proteome of the left ventricle (LV) and isolated ventricular cardiomyocytes (CM) and the phosphoproteome of CM (CM-P) from *Mrgd* ^+/+^ (WT) and *Mrgd* ^−/−^ (KO) mice. The proteomic workflow used in this study is represented in Figure 1A. Since we used WT as the reference dataset (KO *vs.* WT) in this study, up-regulations mean more abundant proteins or phosphorylation sites in KO mice while down-regulations mean less abundant features in KO mice. We identified a total of 3523 proteins. Out of 1285 quantified proteins in the LV, 135 were up-regulated and 130 down-regulated (Figure 1B) while 239 proteins were up-regulated and 115 down-regulated out of 1328 quantified proteins in CM (Figure 1C). Regarding the CM-P, 503 phosphorylation sites were quantified (class I), including 47 phosphosites up-regulated and 33 down-regulated (Figure 1D). The PCA analysis (Figure 1E-G) showed a clear separation between the two experimental groups confirming the impact of the *Mrgd* gene deletion in the cardiac proteome (Figure 1E-F) and phosphoproteome (Figure 1G) of KO mice.

**Figure 1.**
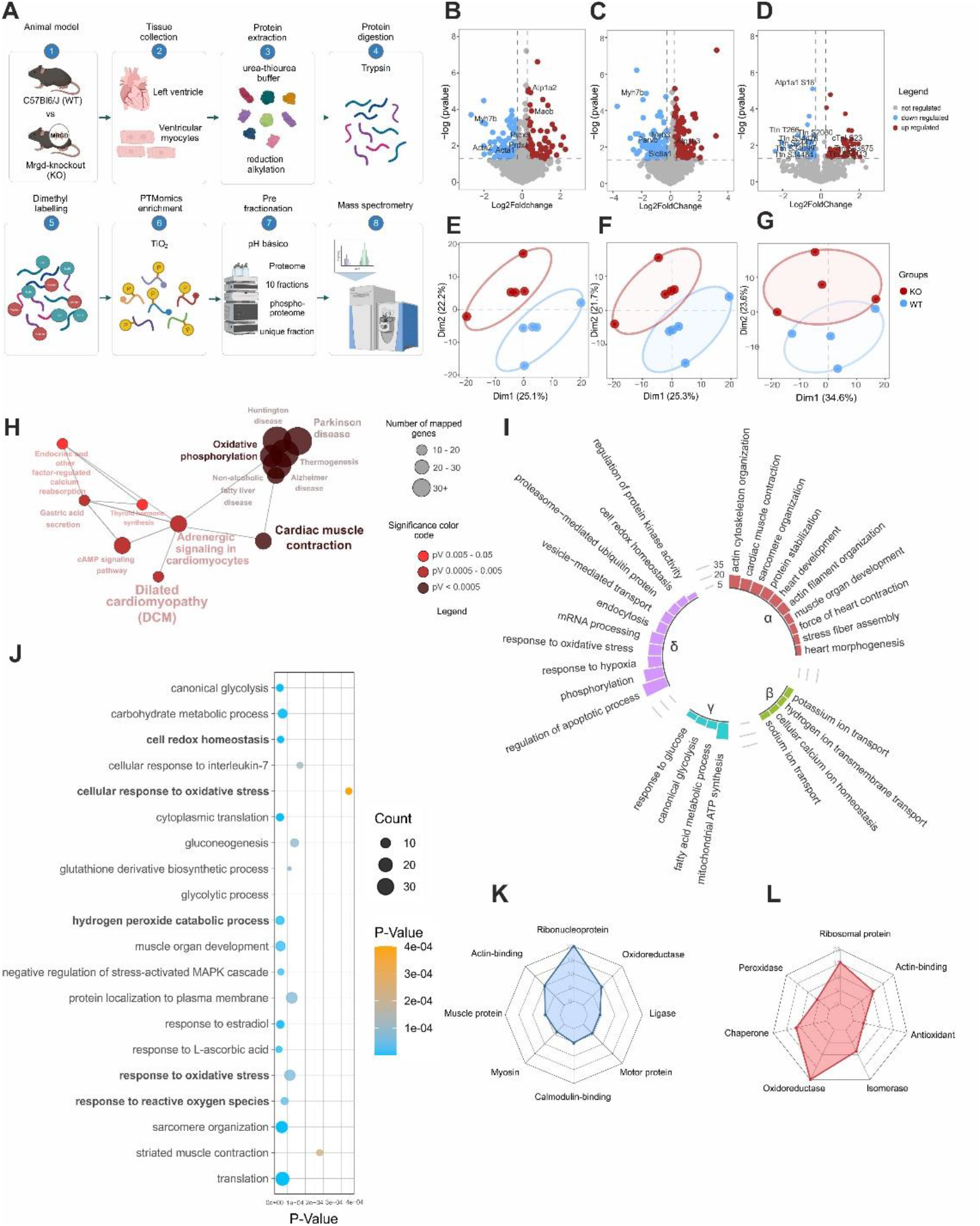
Quantitative proteomics and phosphoproteomics of dilated cardiomyopathy induced by the *Mrgd* gene deletion. (A) Proteomics and phosphoproteomics experimental design. The isolated LV and CM were derived from WT and KO mice (n= 5). Volcano plot of the proteomic data from LV (B) and CM (C). Colored points are regulated proteins with *p*-value ≤ 0.05 and log2 fold-change (log-FC) > 0.26 for up-regulated features, and *p*-value < 0.05 and log-FC < −0.26 for down-regulated features. Gray points represent non-regulated proteins. The *p*-values were determined by one-way ANOVA. Volcano plot of the phosphoproteomic data from CM (D). Colored points are regulated phosphopeptides with *p*-value ≤ 0.05 and log-FC > 0.26 or log-FC < −0.26. Gray points are non-regulated proteins. The *p*-values were determined by one-way ANOVA. Principal Component Analysis (PCA) of proteomics data from LV (E) and CM (F). PCA of phosphoproteomic data from CM-P (G). The KEGG pathways interactome regarding all regulated proteins and phosphoproteins. Each node represents a KEGG pathway, and the edges represent relationships between each term (H). Biological process (BP) terms enriched for the regulated cardiac proteome of MrgD-KO mice. Bar height represents the number of regulated proteins in each GO term. α-δ in (I) represents GO terms related to heart maintenance (α), ion homeostasis (β), energetic metabolism (γ), and oxidative stress (δ). Redundant terms were excluded to present this simplified GO figure but only terms with *p*-value < 0.05 are shown. The detailed GO terms show the top 20 terms with higher significance. The number of regulated proteins in each term (node) is represented in the node size and level of significance is shown in the color key (J). The top 8 molecular function terms using the down-regulated proteins are represented in the spider chart, which shows the number of proteins regulated in each term (K). The top 7 molecular function terms using the up-regulated proteins are represented in the spider chart, which shows the number of proteins regulated in each term (L). Abbreviations: CM, cardiomyocytes; CM-P, cardiomyocyte phosphoproteome; KO, MrgD-KO (*Mrgd* ^−/−^) mice; LV, left ventricle; WT, wild-type (*Mrgd* ^+/+^) mice.

### Gene ontology and pathway analysis associated with MrgD receptor deletion

The regulated features from KO mice were associated with the following KEGG terms: “dilated cardiomyopathy”, “cardiac muscle contraction” and “oxidative phosphorylation”, among others (Figure 1H). The simplified gene ontology (GO) enriched terms (Figure 1I) were involved in heart morphogenesis and cardiac remodeling (panel α), ion transport and homeostasis (panel β), energetic balance (panel γ), oxidative stress (panel δ). A detailed version of the GO showing the top 20 terms associated with the regulated proteins and phosphorylation sites owing to the *Mrgd* gene deletion can be visualized (Figure 1J). Notably, regulated datasets relate to “response to reactive oxygen species”, “response to oxidative stress”, and “hydrogen peroxide catabolic process”. The down-regulated proteins were mainly related to “actin-binding” and “ribonucleoprotein” (Figure 1K) while the up-regulated proteins were associated with “oxidoreductase” (Figure 1L). Thus, the KO cardiac proteome is consistent with a cell redox dysregulation, which can lead to deleterious effects including structural cardiac remodeling ^21,22^ and impaired excitation-contraction coupling ^23^.

### The cardiac tissue from MrgD-KO mice have increased abundance of monoamine oxidase B leading to enhanced oxidative stress

To explore further the mechanisms involved in the generation of a highly oxidative environment in the KO cardiac tissue, we decided to measure the RAS peptides levels in the plasma, bearing in mind the fact that ang II induces ROS production. The experimental levels of circulating ang II, ang-(1-7) and alamandine were not significantly different between KO and WT mice (Figure 2A). Since, a study showed that the expression of angiotensin-converting enzyme (ACE2), ang-(1-7) receptor MAS and ang II receptor AT1 are not different in WT and KO cardiac tissues ^6^, data suggest that the ang II / AT1 signaling pathway does not account for the potential ROS overproduction and consequent cardiac remodeling. However, we observed an increased abundance of MAO-B in the cardiac tissue of MrgD-KO mice (Figure 2B) that might be involved in the potential ROS production (Figure 1I and J).

**Figure 2.**
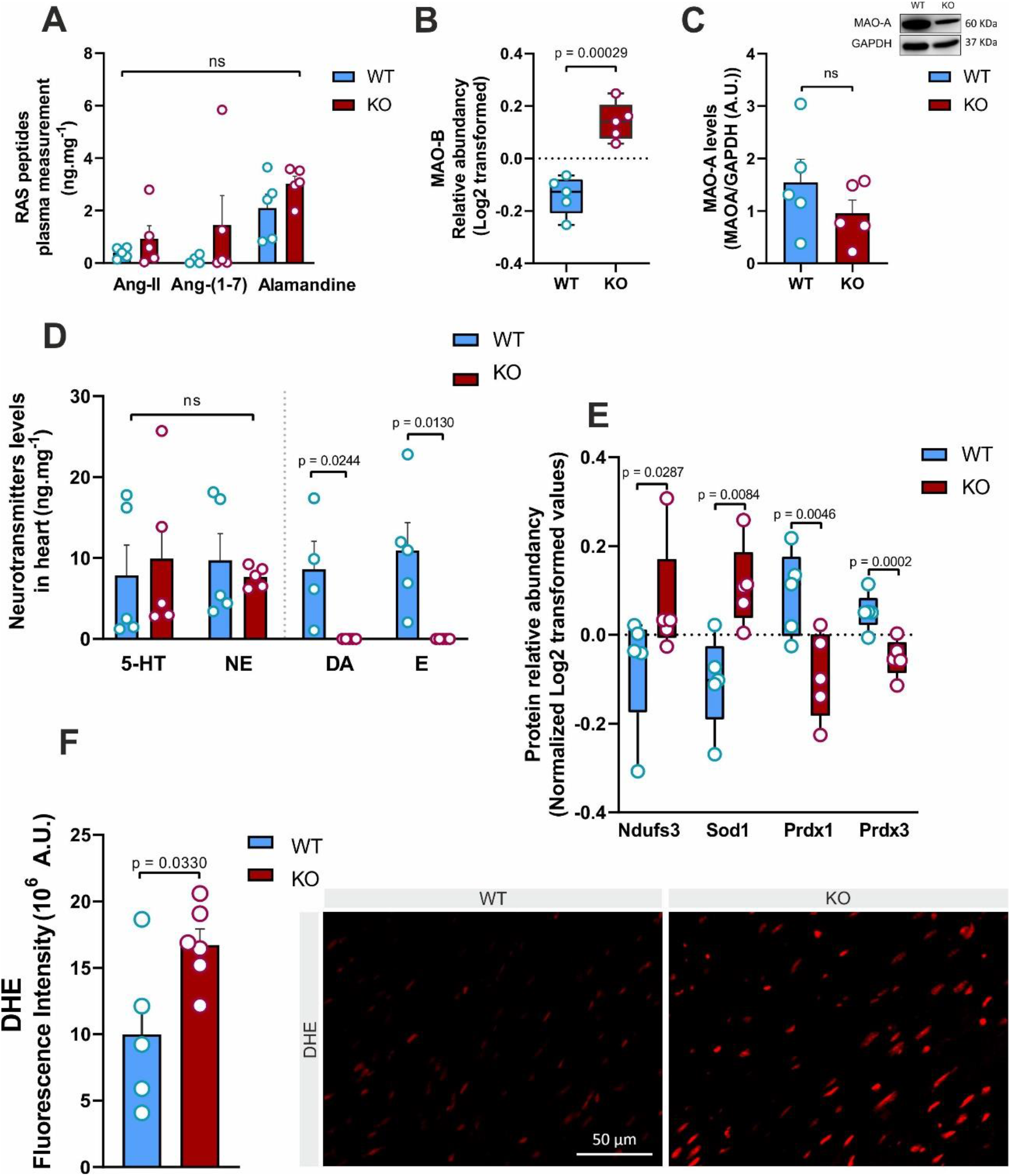
*Mrgd* gene deletion led to increased MAO-B activity and consequent increased generation of reactive oxygen species. (A) Absolute quantification of RAS peptides ang II, ang-(1-7), and alamandine by mass spectrometry (MRM). (B) Relative abundance (log2-transformed value) of MAO-B protein (log2-transformed value) from the left ventricle (LV) proteome. (C) MAO-A protein abundance was similar between the heart of WT and KO mice when measured by western blot. (D) Quantification of the monoamines serotonin (5-HT), norepinephrine (NE), dopamine (DA) and epinephrine (“E”) by high-performance liquid chromatography with fluorescence detector (HPLC-FD). (E) Relative abundance (log2-transformed value) of complex I (Ndufs3), superoxide dismutase type 1 (Sod1), Peroxiredoxin 1 (Prdx1) and (Prdx3) in CM proteome. (F) Measurement of total ROS levels using dihydroethidium (DHE) dye and confocal microscopy. Values are expressed as mean ± SEM and *p*-values were obtained by unpaired t test (two-tailed) in (A; C; D and F). Regulated features were determined as *p*-value ≤ 0.05 and log2 fold-change (log-FC) > 0.26 for up-regulated features and *p*-value < 0.05 and log-FC < −0.26 for down-regulated features. The *p*-values were obtained by one-way ANOVA. Abbreviations: Ang, angiotensin; CM, cardiomyocytes; DA, dopamine; DHE, dihydroethidium; E, epinephrine; KO, MrgD-KO (*Mrgd* ^−/−^) mice; MAO, monoamine oxidase; MRM, multiple reaction monitoring; ROS, reactive oxygen species; WT, wild-type (*Mrgd* ^+/+^) mice.

To investigate whether other MAO isoform would be altered in the failing KO hearts, we quantified by western blotting the abundance of MAO-A, which was not identified by proteomics. As shown in Figure 2C, no significant difference in the cardiac MAO-A was observed between KO and WT mice suggesting that only the abundance of MAO-B is affected by *Mrgd* deletion. Since MAOs are known to degrade monoamines and catecholamines generating hydrogen peroxide (H_2_O_2_), we quantified the cardiac levels of epinephrine (E), dopamine (DA), norepinephrine (NE) and serotonin (5-HT). Interestingly, we observed a depletion of E and DA in the cardiac tissue from KO mice while no significant difference was observed for NE or 5-HT (Figure 2D).

Another potential ROS source in the heart is the complex I subunit (Ndufs3) ^24,25^, which was found up-regulated in KO cardiomyocytes (Figure 2E), suggesting a superoxide overproduction by the electron transport chain (ETC) ^26^. In line with that, the superoxide dismutase (Sod1) was also found up-regulated in KO cardiomyocytes (Figure 2E), suggesting a higher conversion of the superoxide ion (O_2_^−^) into H_2_O_2_ ^27^. Thus, given that H_2_O_2_ is the main product of MAOs and may be produced by Sod1, the abundance of the antioxidant proteins peroxiredoxin 1 (Prdx1) and peroxiredoxin 3 (Prdx3) were explored. Notably, Prdx1 and Prdx3 were found down-regulated in KO mice (Figure 2E), supporting the hypothesis of a highly pro-oxidative environment in the KO cardiac tissue. This was further confirmed by measuring the total amount of ROS by confocal microscopy. ROS levels were significantly higher in the cardiac tissue from KO mice when compared to WT animals (Figure 2F). Thus, we believe that the up-regulation of MAO-B is responsible for the highly pro-oxidative environment observed in the failing KO hearts.

### Cardiomyocytes from MrgD-KO mice have impaired contraction and ionic dynamics

Alterations observed in the proteomic data suggest that the cardiac proteins involved in cardiomyocyte contraction (Figure 1Iα) and ion dynamics (Figure 1Iβ) are dysregulated in KO mice, which prompted us to explore in more details the impact of *Mrgd* gene deletion on these processes. We performed a hierarchical clustering analysis using the regulated proteins and phosphosites associated with “cardiac muscle contraction”, “ion homeostasis”, and “cardiac conduction” KEEG terms. The heat map shows a striking difference between the abundance profiles of the cardiac proteins from KO and WT (Figure 3A). The increase in phosphorylation of the protein kinase A (PKA) at Thr-198 (Figure 3B) and the down-regulation of phospholamban (PLN) and the sodium/calcium exchanger 1, also known as NCX or Slc8a1 (Figure 3C) are consistent with a dysregulated Ca^2+^ homeostasis. To investigate this further, we measured the Ca^2+^ transient in isolated cardiomyocytes. Not surprisingly, a reduced transient amplitude (Figure 3D) was observed in the KO cardiomyocytes. A prolonged decay time (T50) was also observed (Figure 3E), suggesting a reduction in Ca^2+^ reuptake. In line with that, all measured contractile parameters were significant lower in KO cardiomyocytes: contraction-relaxation time - CRT (Figure 3F), contraction time - CT (Figure 3G), relaxation time - RT (Figure 3H), maximum contraction speed - MCS (Figure 3I), maximum relaxation speed - MRS (Figure 3J), and shortening area - SA (Figure 3K). The representative images from the contractile assay are shown (Figure 3L).

**Figure 3.**
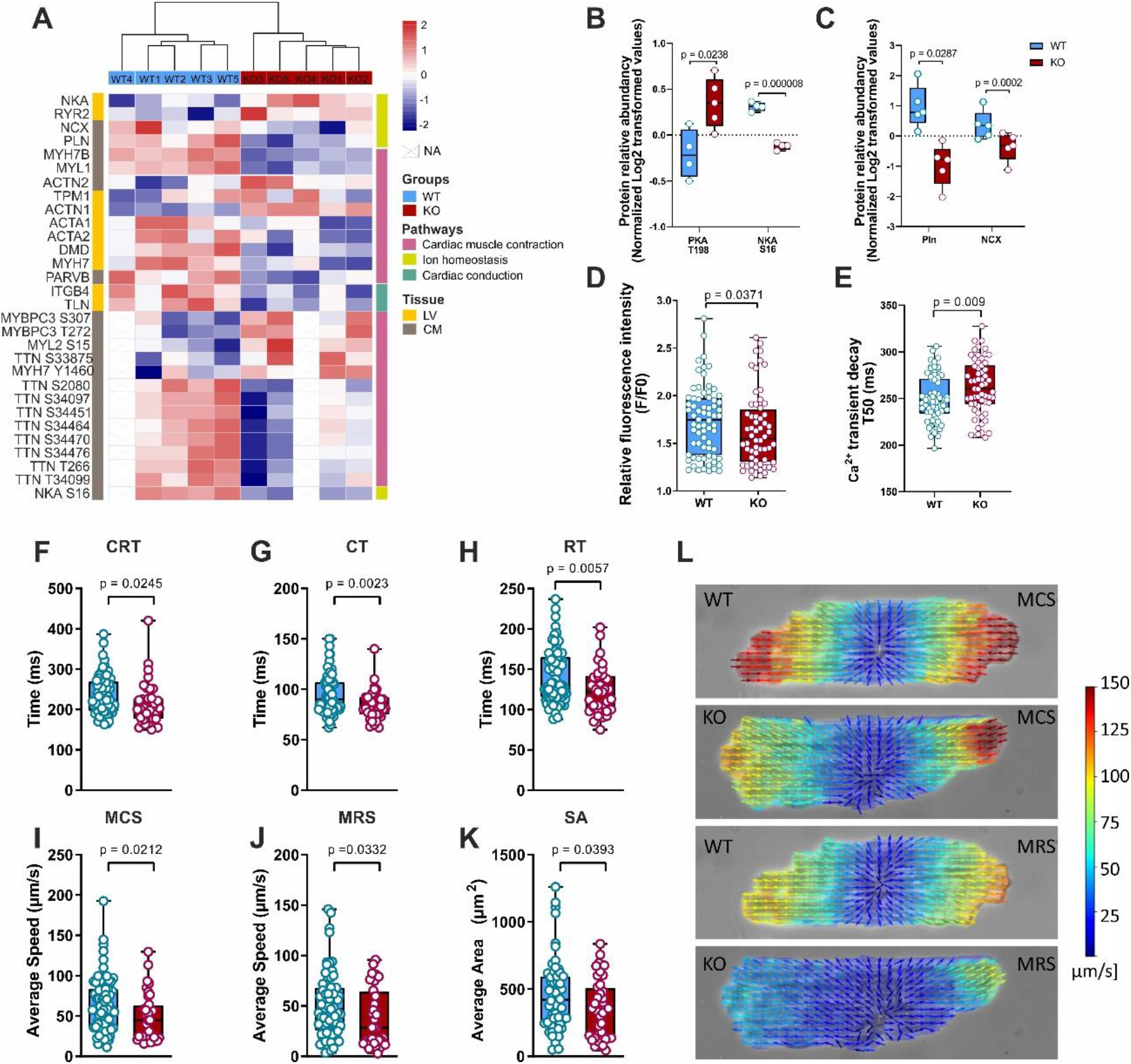
MrgD-KO heart presents an array of molecular features that culminate in dysregulated Ca2+ dynamics and impaired contractions. (A) Heatmap depicting the differentially regulated proteins and phosphosites involved in adrenergic signaling. The heatmap provides a comprehensive view of the altered proteins levels and phosphorylation events in LV and CM from WT and MrgD-KO mice. Mass spectrometry-based relative quantification of (B) PKA phosphorylation at Thr-198 and NKA at Ser-16, and (C) phospholamban (Pln) and sodium/calcium exchanger 1 (Slc8a1/NCX). (D) Ca^2+^ transient amplitude measurement using relative fluorescence intensity (F/F0) (E). Ca^2+^ transient decay (T50, time to 50% decay of the Ca2+ transient)) n=60. Cardiomyocyte (F) contraction-relaxation time (CRT), (G) contraction time (CT), (H) relaxation time (RT), (I) maximum contraction speed (MCS), (J) maximum relaxation speed (MRS), and (K) shortening area (SA). (L) Representative images from cardiomyocytes contraction-relaxion measurements. Regulated proteins were determined as *p*-value ≤ 0.05 and log2 fold-change (log-FC) > 0.26 for up-regulated features and *p*-value < 0.05 and log-FC < −0.26 for down-regulated features. The *p*-values were obtained by one-way ANOVA. Abbreviations: AP, action potential; CM; cardiomyocytes; KO, MrgD-KO (*Mrgd* ^−/−^); LV, left ventricle; RMP, resting membrane potential; WT, wild-type (*Mrgd* ^+/+^) mice.

We also observed an up-regulation of the sodium/potassium-transporting ATPase (Atp1a1 or NKA) in the proteome data (Figure 3A), which was confirmed by western blot (Figure S1A), which might suggest an increased NKA / Atp1a1 activity in KO mice. However, the observed dephosphorylation of NKA / Atp1a1 at Ser-16 (Figure 3B) is consistent with its inhibition and a lower Na^+^ affinity ^28^. The activity of this Na/K pump was indeed reduced in KO cardiomyocytes (Figure S1B). The observed reduction in NKA activity may indicate an increased intracellular concentration of Na^+^, reducing its resting membrane potential (RMP). To test this hypothesis, we measured the RMP from cardiac myocytes by electrophysiology. Indeed, the RMP was more positive in cardiomyocytes derived from the cardiac tissue of KO mice (Figure S1C). No significant difference was observed in the action potential (AP) amplitude (Figure S1C) or in the rise time (Figure S1D), though the AP50 time (50% of repolarization) was lower in KO cardiomyocytes (Figure S1D). The observed lower AP50 time is consistent with the prolonged Ca^2+^ decay in KO (Figure 3E).

### MrgD-KO cardiac myocytes undergo post-translational changes

In addition to the dysregulation in the ion dynamics, several proteins involved in “dilated cardiomyopathy”, “cardiac muscle contraction”, “adrenergic signaling in cardiomyocytes” and “focal adhesion” were dysregulated in KO (Figure S2A). The abundance of actin alpha 1 (Acta1) and myosin heavy chain 7b (Myh7b) were down-regulated in the cardiac tissue of KO mice (Figure S2B), suggesting a decrease in the abundance of thin and thick sarcomere filament proteins. In line with that, the normalized heart weight of KO was lighter than WT (Figure S2C). We measured the cardiac fibers, but no difference in its diameter or structure was observed (Figure S2D). Similarly, the assessment of the cardiac fibers by electronic microscopy did not show any pronounced impairment in their structure (Figure S2Eα-β). However, the mitochondrial shape and the inner mitochondrial membrane seem to be altered when comparing the cardiac tissue from KO (Figure S2Eγ) and WT (Figure S2Eδ). Even though we noticed structural and density changes in the mitochondria from the KO cardiac tissue, the ATP production (in LV and CM) was not impacted, suggesting a normal mitochondrial activity (Figure S2F).

In line with the down-regulation of collagen type IV and VI (Figure S2B), we observed a reduction in the total collagen deposition (Figure S2G) in KO heart tissue.

The cardiac troponin I (Tnni3/cTnI) was found more phosphorylated at Ser-23 in (Figure S2H), and titin (Ttn) was found mainly dephosphorylated, at Thr-266 (disk Z portion), Ser-2080 (Band I portion), and Thr-34099, Ser-34464, Ser-34470, and Ser-34476 (Band M portion) in KO cardiomyocytes (Figure S2H).

### MAO-B inhibition reverts the contractile dysfunction observed in MrgD-KO cardiomyocytes

To investigate if the observed contractile dysfunction of KO cardiomyocytes was associated with MAO-induced ROS production due to the degradation of cardiac monoamines, we used selective MAOs inhibitors and performed the same contractile assay. Isolated cardiomyocytes were treated with Clorgyline and Pargyline, which are selective inhibitors of MAO-A and MAO-B respectively.

In cardiomyocytes isolated from WT mice, the inhibition of MAO-A and MAO-B resulted in a reduction of the CRT (Figure 4A), CT (Figure 4B) and RT (Figure 4C), with no significant difference observed for KO myocytes. However, in contrast to the previous parameters, MAO-B inhibition resulted in increased MCS (Figure 4D), MRS (Figure 4E), and SA (Figure 4F) in KO cardiomyocytes with no difference observed in WT myocytes.

**Figure 4.**
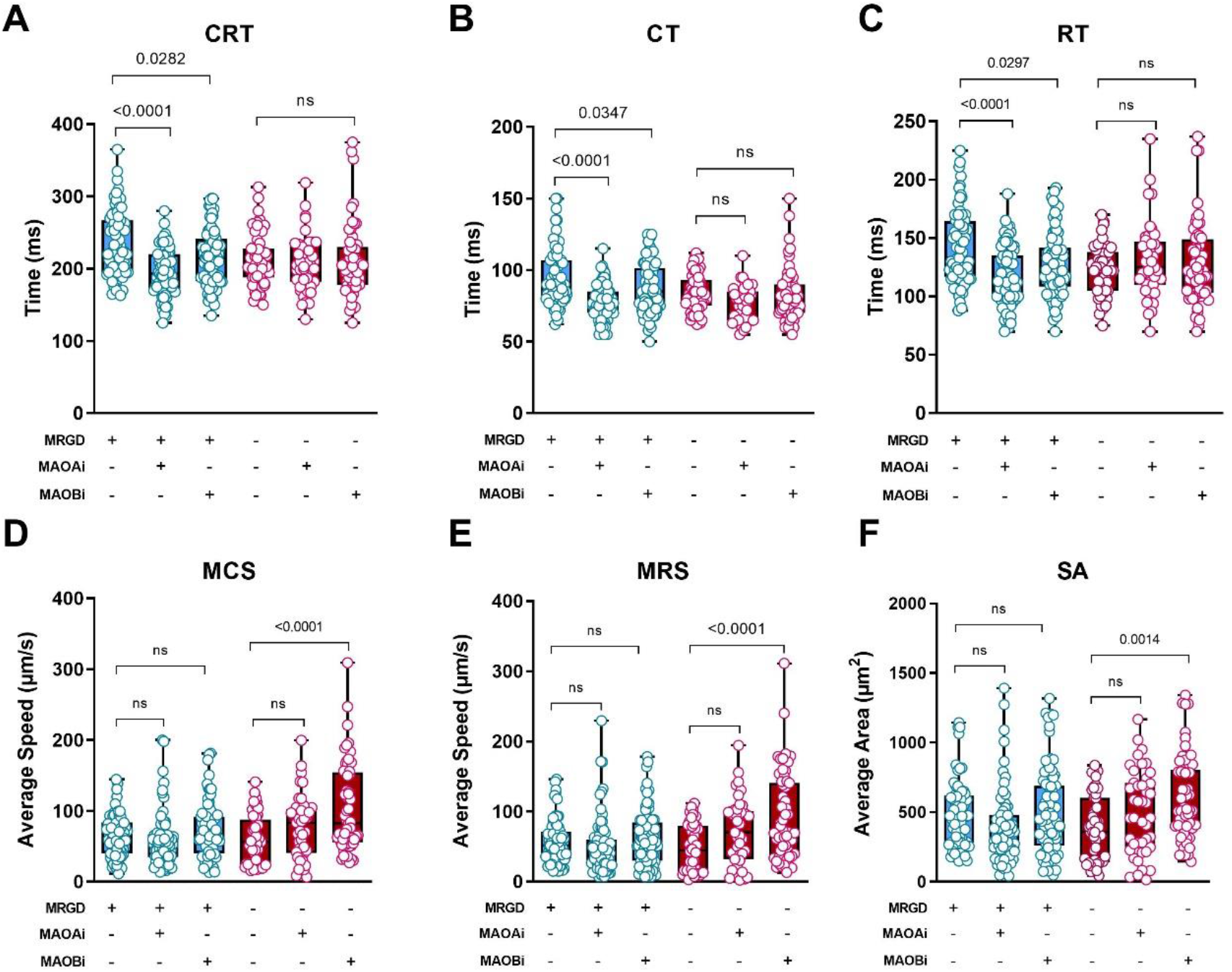
Inhibition of MAO-B but not MAO-A ameliorates the contraction impairment observed in MrgD-KO cardiomyocytes. (A) Contraction-relaxation time (CRT), (B) contraction time (CT), (C) relaxation time (RT), (D) maximum contraction speed (MCS), (E) maximum relaxation speed (MRS), and (F) shortening area (SA) measurements in cardiomyocytes isolated from WT and KO heart in the absence (-) or presence (+) of MAO-A or MAO-B inhibition.

## DISCUSSION

In the study conducted by Oliveira et al.^6^, it was shown that mice lacking the MrgD receptor have a deleterious DCM phenotype, represented by significant reduction in both fractional shortening and ejection fraction, coupled with a remarkable increase in the end-diastolic and end-systolic volumes, and LV asynchronous strain, culminating in a decreased cardiac output. Here, we sought to study the cardiac proteome and phosphoproteome of KO mice in an attempt to unveil the molecular grounds of the DCM associated with the *Mrgd* gene deletion.

### Increased MAO-B activity and ROS production in MrgD-KO mice impair cardiomyocytes contraction

The proteomic data showed a remarkable alteration in pathways often described in other DCM models ^29,30^, such as cardiomyocyte contraction, adrenergic signaling in cardiomyocytes, and dilated cardiomyopathy (Figure 1). We observed proteins related to cell redox and oxidative stress signaling impacted by the *Mrgd* gene deletion (Figure 2). The RAS components are involved in redox homeostasis ^31–33^. However, the data reported here do not support RAS as the source for ROS generation in the KO group (Figure 2A). Instead, the observed up-regulation of MAO-B (Figure 2B) may play a role, potentially contributing to the degradation of DA and E (Figure 2D) and subsequent formation of ROS in the cardiac tissue of KO mice.

The monoamine oxidases (MAOs) are enzymes located within the outer mitochondrial membrane ^34^, that catabolize monoamines generating ROS and other products ^35–37^. Concomitant with the up-regulation of MAO-B (Figure 2B), we observed a more disorganized mitochondrial cristae architecture (Figure S2Eδ), without alteration in ATP production (Figure S2F) in the KO cardiac tissue, which can be an indicative of mitochondrial stress. Moreover, higher expression and increased activity of MAOs have been linked to heart failure, including in humans ^36,38,39^. In addition, the increase in DA catabolism was found in failing hearts, and cardiomyocytes from MAO-B knockout produce less ROS in cardiac tissues ^40,34^. Similarly, the treatment with alamandine (MrgD endogenous agonist) reduced ROS production in transverse aortic constriction ^32,41^. Thus, the antioxidative effect promoted by alamandine / MrgD signaling in the heart seems to regulate MAO-B activity.

Other studies have shown that increased ROS production due DA degradation by MAOs has a notable impact in Ca^2+^ ion dynamics, reducing NKA activity, and leading to a significant impact on cellular excitability ^34,36,42–48^. We observed a reduction in the Ca^2+^ transient accompanied by its reduced reuptake (Figure 3D and 3E), cardiomyocyte contractile dysfunction (Figure 3F-K), and reduced NKA (Atp1a1) activity (Figure S1B).

### Increased ROS in MrgD-KO mice seems to be important for cardiac tension alteration

The increased redox signaling potentially trigged by MAO-B observed in KO cardiac tissue (Figure 2) are commonly related to oxidizing myofilament proteins ^49,50^, contributing to a higher degradation rate of myofilaments such as α-actin and myosin ^51^. In agreement with that, both Acta1/2 and myosin heavy chain 7 (Myh7b) were found down-regulated in KO cardiac tissue (Figure S2B) which could explain, at least partially, the cardiomyopathy observed in KO mice ^52,53^.

The oxidative stress can also disrupt the transmission of calcium-dependent contractile responses ^54–56^. A highly oxidative environment seems to modulate phosphorylation in sarcomere proteins altering its functioning ^57–59^. Treatment of cardiomyocytes with H_2_O_2_ induce the phosphorylation of cardiac troponin I (cTnI/Tnni3) at Ser-23, leading to a decrease in myofilament Ca^2+^ sensitivity, contributing to lower isometric tension and shortening in cardiomyocytes ^57,60–62^. Similarly, we observed a phosphorylation of cTnI at Ser-23 (Figure S2H) and altered cardiomyocyte contraction time, speed and shortening in KO (Figure 3K-L). Likewise, phosphorylation-mediated modifications in titin (Ttn) can induce alterations in its stiffness and passive tension properties. The hypo- phosphorylation of Ttn was found in a cardiac hypertrophy with increased stiffness ^58,59,63^. Similarly, the higher oxidative environment (Figure 2F) together with the observed hypo-phosphorylation of titin (Figure S2H) suggests a potential alteration in the KO cardiac stiffness ^64^, which is consistent with the decreased longitudinal endocardial velocity and circumferential displacement previously reported for KO mice ^6^.

### The role of MAO-B in the observed MrgD-KO contractile dysfunction

Cardiomyocytes isolated from the DCM hearts (KO) treated with the MAO-B selective inhibitor showed contraction improvement, represented by an increase in the maximum contraction and relaxation time (Figure 4D-E) and shortening area (Figure 4F). Similarly, the deficiency of *Maob* gene in mice induced a protective effect when the heart was submitted to a chronic pressure overload, and reverted the contractile injury promoted by ischemia/reperfusion ^34,40^. It is worthy to highlight that the treatment of cardiomyocytes derived from the hypertensive rat TGR (mREN2)27 with alamandine promoted the opposite effects, enhancing contractility, promoting the phosphorylation of PLN on Thr-17 and a higher Ca^2+^ transient and accelerated Ca^2+^ reuptake ^65^. In addition, the alamandine treatment of failing hearts due to myocardial infarction reduced oxidative stress and reverted the cardiac damage caused by ischemia/reperfusion, increasing ejection fraction and fractional shortening ^66^.

## CONCLUSIONS

The molecular mechanisms underlying DCM promoted by *Mrgd* gene deletion involve the up-regulation of MAO-B, leading to increased ROS generation, disrupted calcium handling, impaired contractility, derangements in cardiac structure and function, and alterations in ionic dynamics. The observed improvement in the contractility of MrgD-KO cardiomyocytes after MAO-B inhibition indicates that this enzyme is involved in the observed DCM associated with the lack of MrgD. This new understanding of the impact of MrgD in MAO-B activity opens new avenues for potential therapeutic interventions to mitigate cardiac dysfunctions associated with altered monoamine metabolism. The study provides important insights into the pathogenesis of DCM and suggests potential new therapeutic targets for managing failing hearts.

